# Individual-based modeling of direct and indirect competition by the glass-box artificial intelligence

**DOI:** 10.1101/2025.08.23.671922

**Authors:** Vyacheslav L. Kalmykov, Lev V. Kalmykov

## Abstract

The main drawback of traditional ecological models is the inability to model local interactions between individuals. As a consequence, it was impossible to mathematically model individual-based mechanisms of direct and indirect competition, which hampers the understanding of ecological dynamics. To overcome this problem, we developed a fully transparent mechanistic analog of the classical Lotka-Volterra phenomenological model based on explicitly explainable artificial intelligence (AI) of the glass-box type. In this paper, on an individual-based model, we present (i) a rigorous mechanism of inverted competitive exclusion where, all else being equal, the weaker species in direct competition defeats the stronger species through indirect exploitative competition and (ii) a rigorous mechanism of coexistence of complete competitors resulting from a combination of accounting for microhabitat regeneration, moderate reproduction, a certain habitat size and boundary conditions. Both presented mechanisms go beyond the classical formulations of the competitive exclusion principle, as in our model two species with different competitiveness compete for a single limiting resource in one homogeneous bounded ecosystem. We have shown that individual-based modeling of direct and indirect competition mechanisms allows for a more detailed understanding of the causal mechanisms of interspecific competition. Such understanding is necessary to refine formulations of the competitive exclusion principle, to search for mechanisms of competitive coexistence, to predict ecosystem behavior, and to develop conservation strategies. Our approach allows us to study discrete systems with fundamentally discrete mathematics, which opens up new opportunities for achieving accuracy, clarity, rigor, and explainability in studies of complex systems.

**Highlights:**

- Mechanisms of direct and indirect competition in one individual-based model
- A mechanism of indirect exploitative competition by which a weak competitor wins
- A mechanism of coexistence of complete competitors
- Cellular automata for the eXplicitly eXplainable AI (XXAI) of the glass-box type
- ODD protocol of the model and a source code of the Monte Carlo experiment in C++

## Introduction

The main problem with ecological modeling is that most ecologists use a phenomenological mathematical apparatus to model complex systems. In other words, these models are “black box” types [1]. Traditionally, most ecologists “studied competition by asking if an increase in the density of one species leads to a decrease in the density of another, without asking how this might occur” [2]. The opacity of black box models has made it impossible to study local, individual-based mechanisms of interspecific competition directly [1–4]. Consequently, modeling the mechanisms of direct and indirect interspecific competition at the individual level has largely been unavailable to date. However, modeling indirect competition is essential for understanding how species influence each other indirectly through shared resources or environmental modifications, resulting in complex interactions beyond a direct contact. To solve this problem, we model direct and indirect competition by the explicitly explainable artificial intelligence (AI) of the glass-box type [5]. We use here the term “glass-box” because it more clearly than its synonym “white-box” characterizes the accessibility to control and analysis of the internal mechanisms of the object under study. We have previously used this method to thoroughly test the principle of competitive exclusion and to investigate many aspects of ecosystem dynamics and mechanisms of interspecific competition [4, 6–9]. A model of S-shaped and double-S-shaped single-species population growth [10] is the basis for our models. In [9], we used this model to investigate the catastrophic death of a single population that occurs when the duration of resource regeneration increases.

In this study, we investigate in detail the role of direct and indirect competitive relationships in mechanisms of competitive exclusion and competitive coexistence. We are most interested in the mechanism of interspecific competition when a weak complete competitor, other things being equal, displaces a strong one in a closed, initially homogeneous habitat. This mechanism required a deeper investigation of the underlying causal mechanisms. A mechanism in which a weak complete competitor can win was first discovered and presented by us using a minimalist model [8]. In this minimalistic model, we simulated aggressive competition between individuals of the strong and the weak species on a one-dimensional, three-cell field with a fixed boundary condition, all other things being equal. Relative competitiveness was defined as competitiveness under conditions of direct competition between individuals for a vacant microhabitat. Individuals reproduce aggressively, competing on a case-by-case basis for microhabitat with a free limiting resource. Aggressive reproduction is modeled by a hexagonal neighborhood. Fig. 1 shows mechanisms of two types of competitive exclusion resulting from direct (Fig. 1A) and indirect (Fig. 1B) competition and a mechanism of competitive coexistence (Fig. 1C).

**Figure 1.**
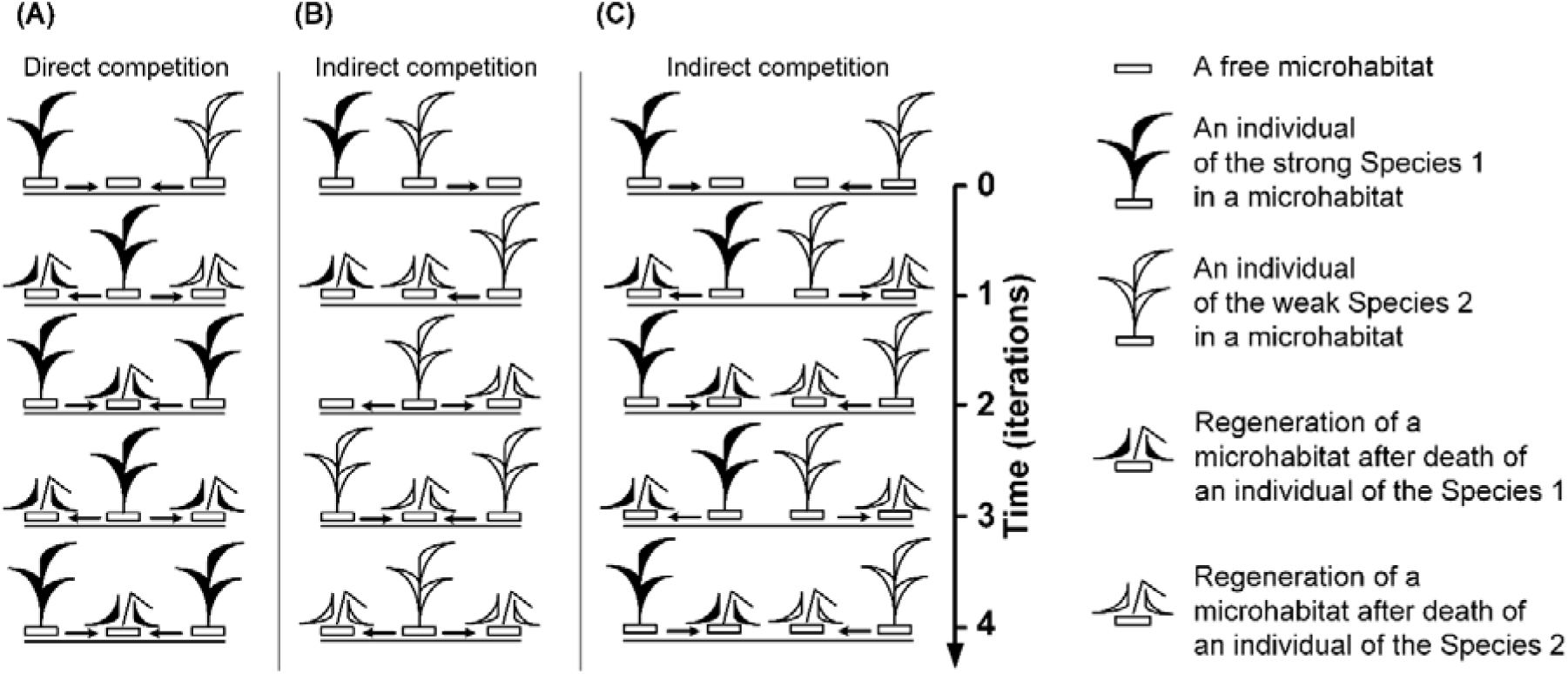
Deterministic individual-based mechanisms with local interactions. One-dimensional field with fixed boundary and hexagonal neighborhood. Arrows mark potential reproduction directions of individuals at the next iteration. (A) Competitive exclusion of the weak Species 2 as a result of the direct conflict for a free microhabitat. (B) Inverted competitive exclusion of the strong Species 1 when the weak Species 2 wins in a result of indirect exploitative competition. (C) Coexistence of the both competing species.

Figure 1A shows the classical competitive exclusion, when the strong Species1 wins in a direct conflict of interest. Figure 1B shows the case of inverted competitive exclusion - the victory of the weak species as a result of indirect exploitative competition. An individual of the weak species physically blocks an individual of the strong species in the “corner of the habitat” from gaining access to the only available resource for propagation. At the same time, the weak species has an access to the free resource while avoiding a direct local competitive conflict. The only difference between these two cases is the different initial placement of individuals in space. Figure 1C shows an example of an individual-based mechanism of coexistence of complete competitors.

The aim of this study is to demonstrate, based on an individual model, the role of direct and indirect interactions between species in competitive mechanisms. The greatest attention is paid to identifying cases of inverted competitive exclusion in a two-dimensional habitat model.

Additionally, we examine how the inverted competitive exclusion and coexistence of complete competitors found here relate to our formulations of the principle of competitive exclusion [6, 8].

## 1. Methods

### 1.1. Explicitly explainable AI of the glass-box type

Understanding the mechanisms underlying natural systems requires identifying causal interactions [11]. We use our model, which is based on the axioms of general ecosystem theory, to conduct an individual-based investigation of the role of direct and indirect interactions between competing species using automatic causal inference. [6–8]. The model is a logical, deterministic cellular automaton. It simulates cause-and-effect mechanisms of competition by explicitly considering the local interactions of competing individuals. This individually-based mechanistic model is constructed using an explainable artificial intelligence method of the glass box type. [4, 5, 7, 10]. Automatic causal inference is realized as a general network inference on the whole field of the cellular automaton. The cellular automaton neighborhood ensures the integrity of the general solution by gluing the local logics of state changes of individual cells into a single whole. The evolution of such a model provides transparent results that allow direct identification of the deterministic causal mechanisms of competition. The logical rules of the cellular automaton are applied to all microobjects (cells), to their corresponding mesoobjects (neighborhoods), thus covering the entire macro-level (cellular automaton field) of the modeled complex system at each time iteration. Such models possess discreteness, integrity, nonlinearity, emergent, parallelism, operational and semantic transparency, accuracy of solutions without errors, rounding and artifacts.

**The discreteness** of the deterministic logical cellular automaton is due to the fact that it operates within discrete time and space, and uses a finite set of states for each cell of the field.

**The integrity** of the overall solution is ensured by the fact that the cellular automaton neighborhood glues the logic of local changes of states of all cells of the field into a single pattern-solution.

**Nonlinearity** of the cellular automaton dynamics is due to the fact that small changes in initial conditions may lead to large significant changes in the behavior of the system.

**The emergence** of the behavior of a cellular automaton is realized as its self-organization, the result of which cannot be predicted in advance. This result is the final evolutionary pattern of the cellular automaton. A cellular automaton solves a problem by transforming an existing pattern into an updated configuration that serves as a global, logical solution. This solution is implemented as local rules that are applied synchronously to all cells. These local rules are based on the first principles of the corresponding problem domain. The emerging pattern visualizes the transformed local logical calculations.

**The parallelism** of the cellular automaton is related to the fact that the states of all cells of the field are updated synchronously and iteratively.

**Operational transparency** of the model means that it allows to control internal mechanisms at micro-, meso-, and macro-levels at all stages of modeling.

**Semantic transparency of the model** means that it is built using explicit concepts at all levels of model organization. This is achieved by the fact that these explicit concepts are objects of the axiomatic theory of the problem domain. This theory serves as the knowledge base of the cellular automata expert system.

**The accuracy** of solutions without errors, rounding and artifacts is ensured by logical deterministic transitions between a finite set of integer states.

These features, when combined, create cellular automata with rules based on the fundamental principles of their respective subject domains. This makes them a powerful tool for transparent modeling and studying the mechanisms of complex systems.

This method allowed us to identify a number of subtle mechanisms of competition. In a real ecosystem, many such mechanisms may operate simultaneously. This is due to intraspecific diversity and the diversity of real environmental conditions. Each competitive mechanism is sensitive to the conditions in which it operates. However, this does not make the mechanism artificial or diminish its importance. A metaphorical parallel can be drawn. A grain of sand is enough to upset the clock. However, this does not diminish its importance of this mechanism. Similarly, a precise deterministic mechanism of competition, revealed by the automated inference is not an artifact, even though it is potentially influenced by initial conditions.

### 1.2. Modeling interspecific competition

The neighborhood of cellular automaton (Fig. 2A,B) determines the potential ability of individuals to use free resources of the environment to realize their main task - reproduction. The cellular automaton neighborhood of a microhabitat with the living individual defines potential microhabitats for populating offspring of that individual in the next iteration. A free microhabitat contains all resources for a single offspring. The complete discreteness of the individual-based model allows us to distinguish between direct competition (interference), and indirect competition (exploitation). Direct competition (interference) occurs as a direct conflict of interest (Fig. 1A). In this case, individuals of competing species interact directly and simultaneously in the struggle for the same limiting resource. This may be simultaneous aggressive competition for the same resource, allelopathy, or other direct competition. Indirect exploitative competition occurs when individuals of competing species consume a limiting resource while avoiding a direct conflict of interest.

**Figure 2.**
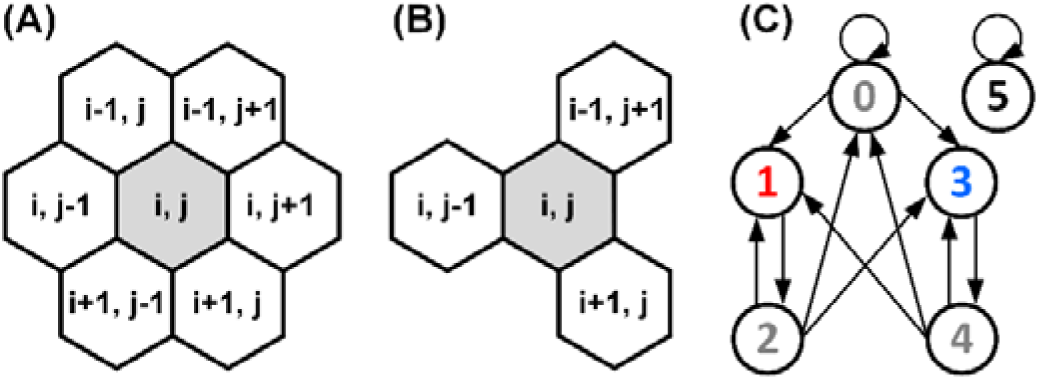
Neighborhoods and a directed graph of the logical transitions of the states of a cell in an ecosystem model with resource competition between two species. (A) The hexagonal neighborhood models aggressive propagation. A central cell and its neighboring cells are defined by the array elements with indices. (B) The tripod neighborhood models moderate propagation. (C) A directed graph of logical transitions between the states of a field cell. Each cell may be in one of the six states: 0 — a free microhabitat which can be occupied by a single individual of any species; 1 — a microhabitat is occupied by a living individual of the Species 1; 2 — a regeneration state of a microhabitat after the death of an individual of the Species 1; 3 — a microhabitat is occupied by a living individual of the Species 2; 4 — a regeneration state of a microhabitat after the death of an individual of the Species 2; 5 — a state of the boundary cell that cannot be occupied.

We investigate interspecific competition using a model of joint asexual colonization of a closed homogeneous habitat. Competitive colonization is initiated by two single individuals of competing species. Individuals of the both competing species consume the same limiting resource, i.e. they are identical consumers. This limiting resource is a free microhabitat containing all the resources necessary for the life cycle of an individual. Individuals of competing species differ in their relative competitiveness under direct simultaneous conflict of interest, i.e., in their functional traits affecting direct competitiveness. Functional traits are morphological, biochemical, physiological, structural, phenological or behavioral characteristics of organisms that influence their fitness [12, 13]. A particular form of adaptive functional traits is allelopathy [14]. We define relative competitiveness as the value of the probability of winning a direct, simultaneous, local conflict of interest between individuals for the same unit of the limiting resource. For clarity, we have shown with arrows the intention of an individual to use the free resource (Fig. 1). The relative direct competitiveness of the strong Species 1 in direct competition in this study is 1, and the same value for the weak Species 2 is 0, i.e. the outcome of direct competition between individuals is fully deterministic. Indirect competition occurs when the offspring of one of the competitors consume a shared, limiting resource without engaging in direct, simultaneous conflict of interest. The separation of direct and indirect interspecific competition provides a transparent mechanistic understanding of the mechanisms involved.

Mechanistic in our case means modeling local, individual-based mechanisms. The model’s semantic transparency is ensured because its rules for controlling the cellular automaton are based on the axioms (first principles) of the general domain theory. The model’s operational transparency is achieved through the cellular automaton’s causal inference, which composes all local causal inferences into a global one. Such models are transparent (glass box) because they are based on discrete time and space, deterministic rules, and explicit consideration of local conditions of individuals’ behavior.

The direct biological prototype of our cellular-automaton model is the competition between individuals of two species of lawn grasses for free microhabitats to accommodate their offspring. These are red fescue (*Festuca rubra*) and meadow bluegrass (*Poa praténsis*), which have numerous cultivars that differ in functional traits. These grasses reproduce vegetatively by creeping underground shoots (rhizomes). The geometry of rhizomes propagation is close to the cellular-automaton neighborhoods we use (Fig. 2A,B), which model the potential geometry of progeny (vegetative seedlings) placement. We model the ecosystem by the entire field of a cellular automaton. We investigate it with fixed nonperiodic and with periodic boundary conditions. A cell containing a deceased individual mimics the microhabitat in a regenerative state. A cell in a regenerating state is unavailable to be populated by a new individual until regeneration is complete. In this way we account for the regeneration niche [7, 15, 16]. We made several assumptions about the duration of cell states. The duration of the ‘occupied’ and ‘regeneration’ states lasts one iteration. The ‘free’ state of a microhabitat is maintained until it is occupied by an offspring of one of the individuals. When an individual dies, its microhabitat goes into the ‘regeneration’ state. After the regeneration state, a microhabitat becomes free if there is no one living individual in its neighborhood. A cell in the regeneration state becomes free or can be occupied immediately after the end of the regeneration state. To implement the boundary conditions, an additional unchanging state of the field cell ‘boundary’ is introduced.

The field cell in the boundary state is always in the same state. A field with a boundary is a field surrounded by an impenetrable “wall”. A field with an impermeable boundary is more consistent with field and laboratory experiments than the situation on a torus, which is modeled by periodic boundary conditions. A periodic boundary conditions of the field is a torus formed by “gluing” its opposite sides together. Figure 2 shows the neighborhoods of cellular automata and the transition graph between the states of the field cells. The descendants of an individual can potentially occupy the nearest free microhabitats in a vegetative way according to the cellular-automata neighborhood. The individual of Species 1 wins in all cases in direct conflict with Species 2 for the only limiting resource for propagation.

To investigate how the initial placement of starting individuals of competing species is affected, we performed Monte Carlo experiments., i.e. a large series of similar experiments under different initial conditions. In each of the 1000 repeated trials, the initial placement of individuals on the field was varied. We have investigated the effect of boundary conditions on the outcome of competition in the 30×30, 31×31, and 32×32 habitats (Figs. 3, 4). Three field sizes were chosen after preliminary tests. In the preliminary tests of smaller and increasingly larger fields under moderate reproduction conditions (tripod neighborhood) and with periodic boundary conditions, we found a periodic repetition of competition patterns through the three field sizes to the fourth. Therefore, to avoid duplication, we limited our study to these three field sizes.

**Figure 3.**
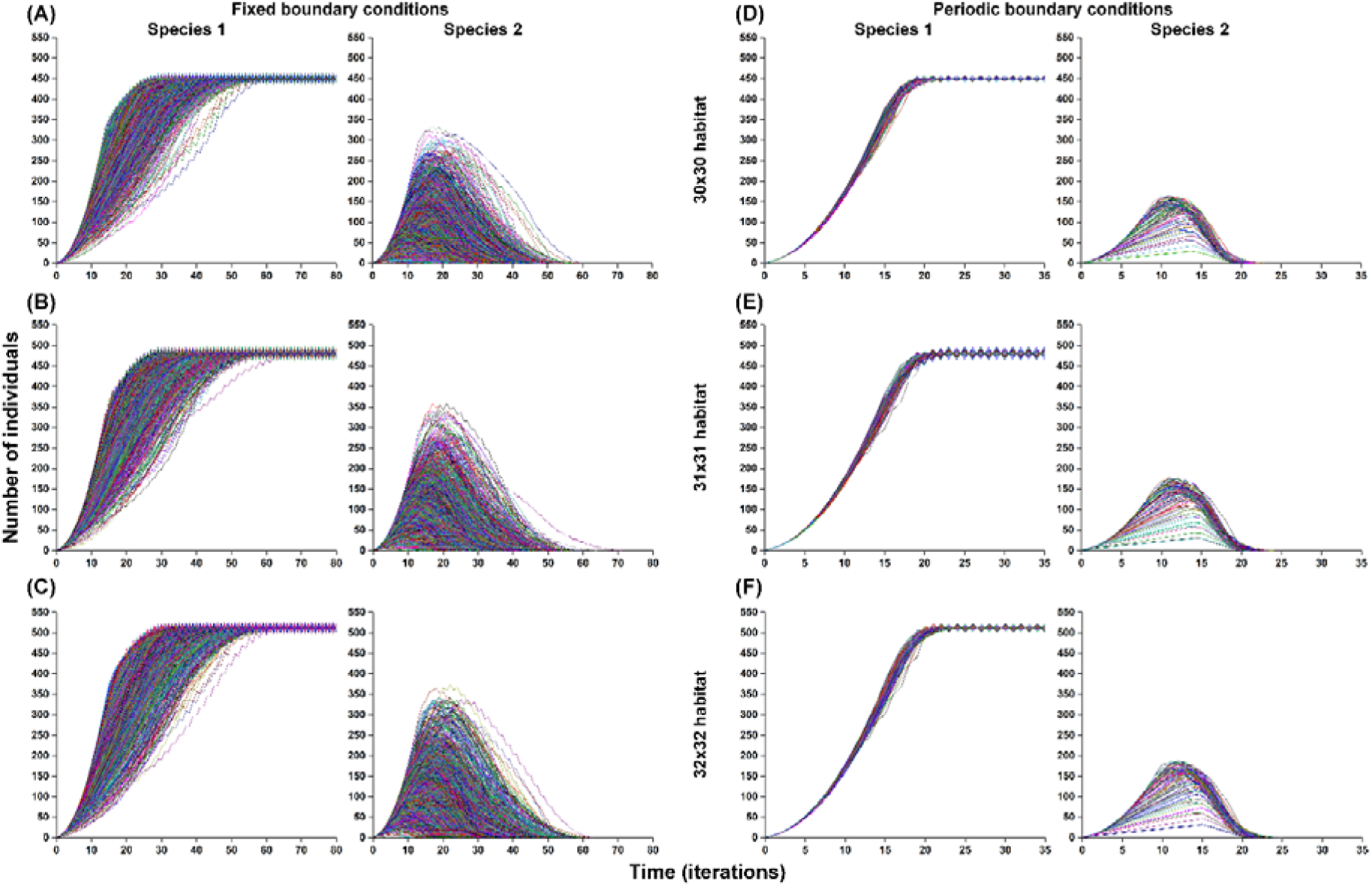
Population dynamics of the model with Monte Carlo simulations (n=1000) when reproduction of the both competing species is aggressive (hexagonal neighborhood). In each of the 1000 repeated trial experiments, the initial placement of individuals of competing species on the lattice was random. Boundary conditions are fixed (A-C) and periodic (D-F). (A-F) Competitive exclusion in all cases. Species 1 wins the Species 2 in all cases. (A) The field size is 32×32 and the habitat size is 30×30. (B) The field size is 33×33 and the habitat size 31×31. (C) The field size is 34×34 and the habitat size is 32×32. (D) The field size is 30×30 and the habitat size 30×30. (E) The field size is 31×31 and the habitat size 31×31. (F) The field size is 32×32 and the habitat size is 32×32.

For a more detailed representation of the mechanism, we also presented single cases of competition on small fields of 3×3 cells patterns on increasingly smaller and increasingly larger fields under tripod neighborhood conditions under periodic boundary conditions. Our model reproduces the competition of complete competitors for one limiting resource in one initially homogeneous habitat. We model the condition of asexual (vegetative) propagation of complete competitors competing for one limiting resource in a homogeneous limited habitat with periodic conditions to exclude both hidden uncertainties that are typical for experimental field studies and numerous mechanisms of competitive coexistence identified earlier [17–19].

Although our models have biological prototypes, they are conceptual, not imitative. Due to their abstract nature, conceptual models are generally significant in understanding the mechanisms of various forms of competition.

In Supplementary I we provide the ‘Overview, Design concepts and Details’ (ODD) protocol of our individual-based model. This protocol aims to create a clear and complete description of the model to facilitate its reproduction.

In Supplementary II we provide a commented C++ source code of the model with Monte Carlo simulation.

## 2. Results

### 2.1. Individual-based modeling of competitive colonization of an ecosystem, taking into account direct and indirect interactions

We investigated population dynamics under different initial conditions of the model (Figs. 3, 4). According to most formulations of the competitive exclusion principle, we should not have found cases of coexistence because “complete competitors cannot coexist” [20]. In the case of colonization of a large two-dimensional habitat by two aggressively reproducible species starting from single individuals, we indeed found no cases of coexistence (Fig. 3). The more competitive Species 1 always displaced Species 2 as a result of the collision of their population waves(Movie S1). However, in the case of moderate reproduction under the same conditions, there were many cases of coexistence (Fig. 4), as well as a victory of the less competitive Species 2. These cases particularly interested us (Movies S2-S4).

**Figure 4.**
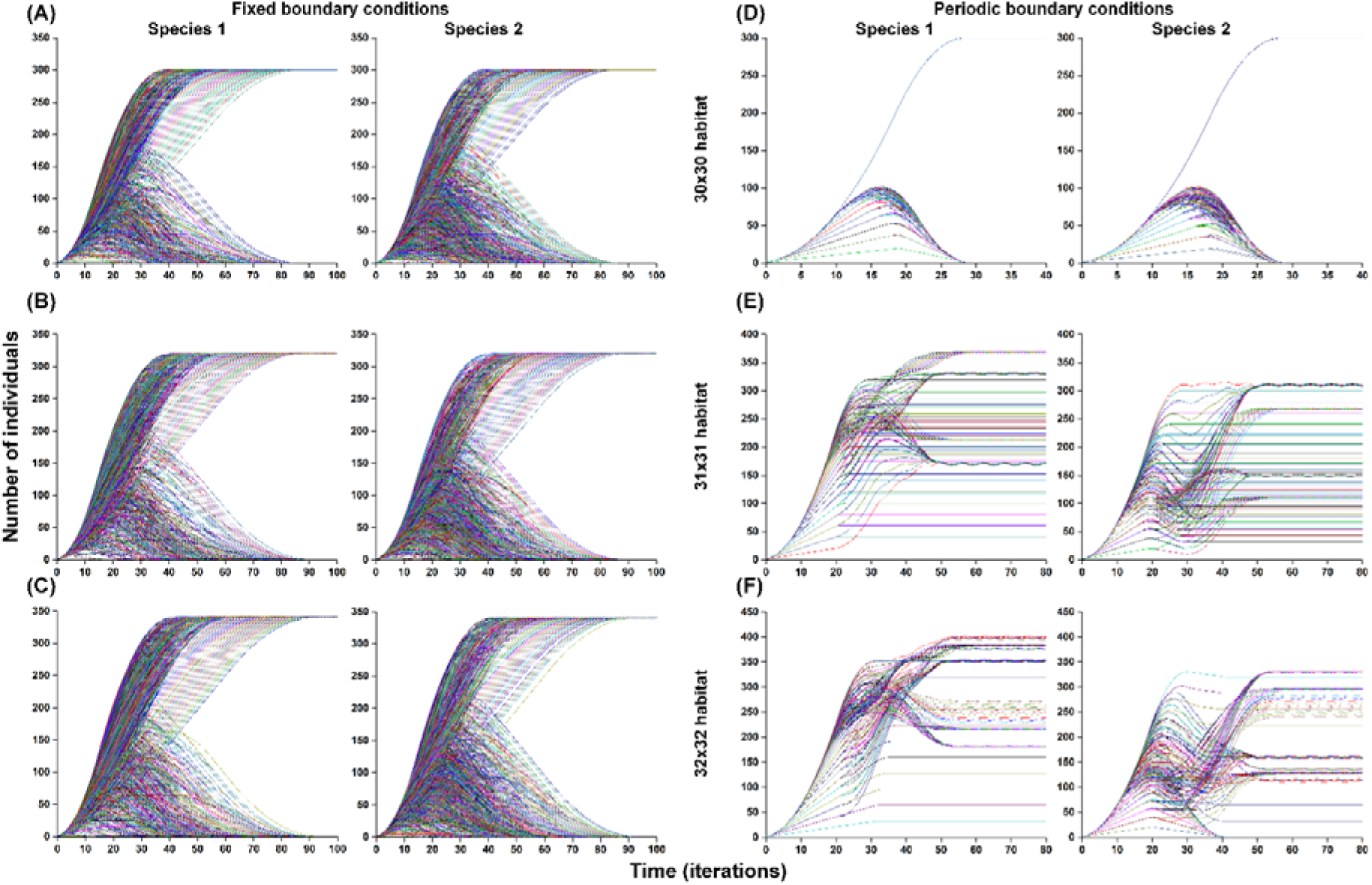
Population dynamics of the model with Monte Carlo simulations (n=1000) when reproduction of the both competing species is moderate (tripod neighborhood). In each of the 1000 repeated trial experiments, the initial placement of individuals of competing species on the lattice was random. Boundary conditions are fixed (A-C) and periodic (D-F). (A) Fixed boundary conditions. The field size is 32×32 and the habitat size is 30×30. The Species 1 wins the Species 2 in 655 cases out of the 1000 cases and the Species 2 wins the Species 1 in 345 cases out of the 1000 cases. (B) Fixed boundary conditions. The field size is 33×33 and the habitat size 31×31. The Species 1 wins the Species 2 in 695 cases out of the 1000 cases and the Species 2 wins the Species 1 in 305 cases out of the 1000 cases. (C) Fixed boundary conditions. The field size is 34×34 and the habitat size is 32×32. The Species 1 wins the Species 2 in 660 cases out of the 1000 cases and the Species 2 wins the Species 1 in 340 cases out of the 1000 cases. (D) Periodic boundary conditions. The habitat size 30×30. The Species 1 wins the Species 2 in 661 cases out of the 1000 cases and the Species 2 wins the Species 1 in 339 cases out of the 1000 cases. (E) Periodic boundary conditions. The habitat size 31×31. The Species 1 and the Species 2 coexist in all 1000 cases. (F) Periodic boundary conditions. The field size is 32×32 and the habitat size is 32×32. The Species 1 wins the Species 2 in 219 cases out of the 1000 cases and the Species 1 and the Species 2 coexist in 781 cases out of the 1000 cases.

We did twelve Monte Carlo simulations, testing different initial conditions for the model. From each of twelve Monte Carlo experiments we obtained 1000 population curves for each species for each specific combination of the cellular automaton neighborhood type, lattice size and boundary condition type (Figs. 3 and 4).

We obtained a variety of mechanisms by applying the Monte Carlo method to the cellular automaton model (see Figures 3 and 4). We analyzed these mechanisms. Fig. 3 shows the results for aggressive propagation of the both competitors, and Fig. 4 shows the results for their moderate propagation. Each Monte Carlo experiment consists of 1000 different trials. The program counts cases of competitive exclusion and coexistence. To compare the population dynamics the results are presented in two different graphs for the Species 1 and the Species 2 respectively (Figs. 3-4).

The individual-based mechanisms of population dynamics of competing species are visible on the cellular automaton field during the model’s evolution. To demonstrate this, we present four videos of computer experiments in the Supplementary materials (Videos S1-S4). Videos S1 and S3 demonstrate the mechanism of the classical competitive exclusion - the strong competitor wins. In both cases, the habitat is 30×30 with periodic boundary conditions. Video S1 shows this mechanism under aggressive reproduction, which includes all cases of exclusion in Fig. 3. Video S3 shows this mechanism under moderate propagation of both species, as shown in Fig. 4.

Most interesting are the cases where the weak Species 2 wins the strong Species 1 (Video S2, corresponding to one of the cases in Fig. 4D). This inverted competitive exclusion is realized in the 30×30 habitat with periodic conditions under moderate propagation of the both species. Video S4, Fig. 1C and Fig. 4 demonstrate mechanisms of coexistence of complete competitors.

We consider two types of boundary conditions - when the field is a torus closed by periodic conditions and when it has a boundary. Periodic conditions are most often studied because this avoids the influence of boundary effects. In the videos shown, the field represents the sweep of the torus (periodic boundary conditions). The population waves in the model propagate on the torus. Species 1 appears behind the corner of Species 2 due to the periodic conditions on the torus (as a result of crossing the boundary of the torus sweep).

The experiments in Figure 3 show that in competitive colonization of a 30×30, 31×31 and 32×32 free habitat with aggressive propagation of the both species. The Species 1 always wins, regardless of the initial placement and boundary conditions. No matter how we place the initial individuals on the field, the strong Species 1 always wins. This result is expected. All population curves in Figure 3 show the classical competitive exclusion, where the strong species wins.

In the experiments presented in Figure 4, we investigated moderate propagation modeled using the tripod neighborhood (Fig. 2B). For a tripod neighborhood, the maximum number of offspring from a single individual is three. For a hexagonal neighborhood, the maximum is six. This can be seen when comparing the two types of the cellular automata neighborhoods in Figures 2A and 2B. Gaps (free microhabitats) are formed in the population waves as a result of propagation in the tripod neighborhood. Both competitors have the same number of offspring, and their traits are identical except for the direct competitiveness. This is important because it eliminates trade-offs and other well-known mechanisms of coexistence. [18].

A weak competitor’s victory is a unique and even paradoxical case. In our model experiments, we observed numerous instances of reverse competitive exclusion that were independent of boundary conditions. (Fig. 4A-D). In all cases of aggressive reproduction, we studied, neither inverted competitive exclusion nor competitive coexistence occurred during colonization of initially homogeneous two-dimensional habitats (Fig. 3).

### 2.2. Cell by cell representation of direct and indirect mechanisms of competition in a 3×3 habitat

For the most visual microscale, cell-by-cell representation of the mechanisms of direct and indirect local competition mechanisms, we present their implementation on small 3×3 microhabitats in two-dimensional fields with fixed boundary conditions (Fig. 5) and periodic boundary conditions (Fig. 6). Only variants under moderate reproduction (tripod neighborhood) were studied, since in this paper we found that on a two-dimensional homogeneous field under aggressive multiplication (hexagonal neighborhood) a strong competitor always wins. Figures 5 and 6 show three different initial patterns. In Fig. 5A, a strong species wins through direct competition. In Fig. 5B, a strong species wins through indirect exploitative competition. In Fig. 5C, the weak Species 2 wins through inverted competitive exclusion.

**Figure 5.**
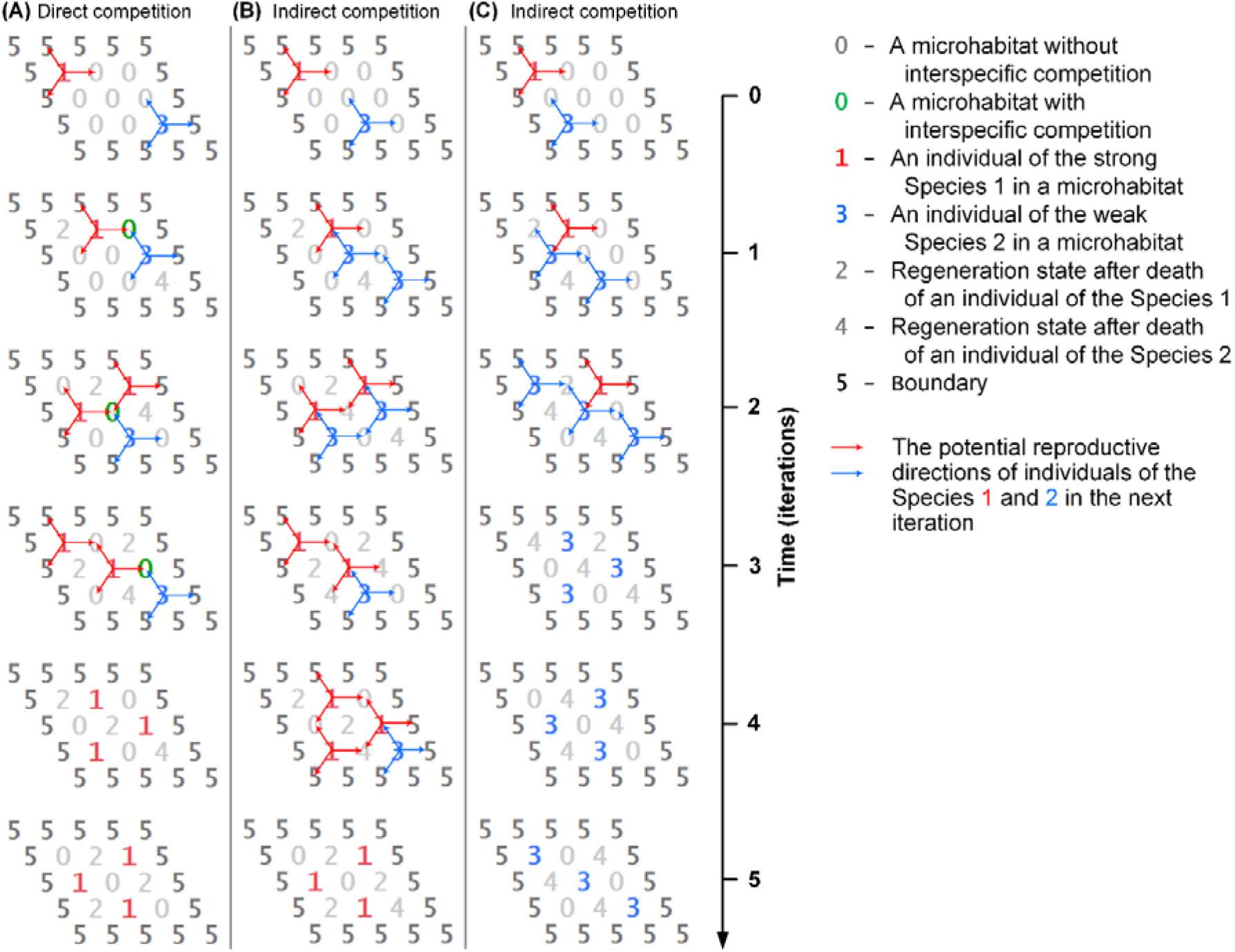
Local mechanisms of interspecific competition on the two-dimensional lattice with the boundary. Habitat size 3×3 with the field size 5×5 (non-periodic boundary conditions). The tripod neighborhood of the cellular automaton simulates moderate vegetative reproduction. (A) Competitive exclusion of the weak Species 2 as a result of direct local conflicts for microhabitats marked in green. (B) Competitive exclusion of the weak Species 2 when the strong Species 1 wins as a result of indirect competition. (C) Inverted competitive exclusion of the strong Species 1 when the weak Species 2 wins as a result of indirect competition.

**Figure 6.**
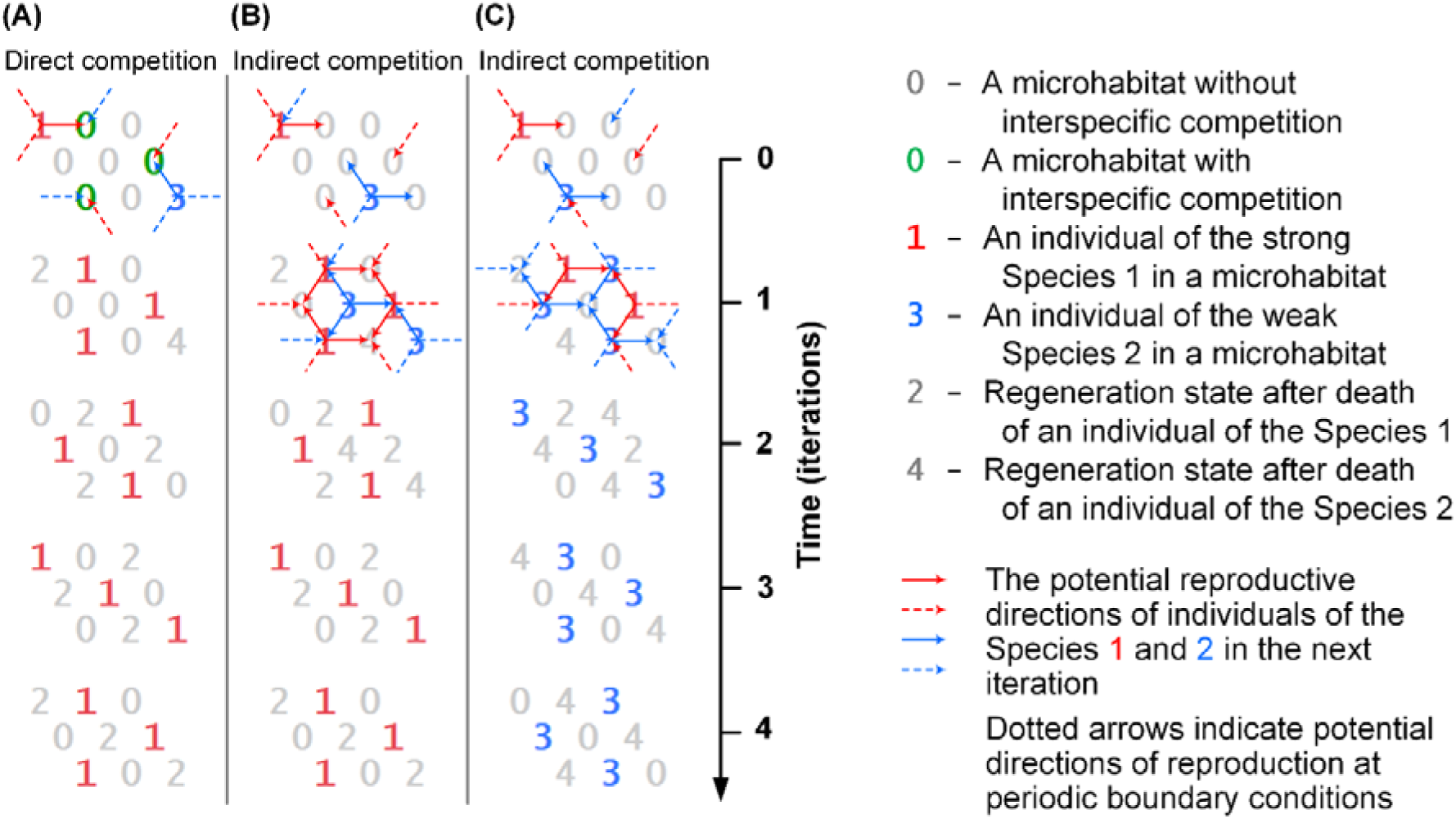
Local mechanisms of interspecific competition on the two-dimensional field closed by periodic conditions. Field and habitat size is 3×3 cells. The tripod neighborhood of the cellular automaton simulates moderate vegetative reproduction. (A) Model evolution results in competitive exclusion of the weak species as a result of three direct local conflicts over microhabitats marked in green. (B) Competitive exclusion of the weak Species 2 when the strong Species 1 wins as a result of indirect competition. (C) Inverted competitive exclusion of the strong Species 1 when the weak Species 2 wins as a result of indirect competition.

Fig. 6 shows: Fig. 6A, the strong species wins through direct competition; Fig. 6B, the strong Species 1 wins through indirect exploitative competition; Fig. 6C, the weak Species 2 wins through inverted competitive exclusion.

## 3. Discussion

Here we show that modeling direct and indirect competitive interactions at the local level allows for a deeper analysis of competition mechanisms (Figures 1, 5, 6; Movies S1-S4). The most striking example is the modeling of inverted competitive exclusion. (Figures 1B, 5C, 6C; Movie S2). It has been demonstrated that this is not an isolated or rare case, but rather a fairly common occurrence. It is important to note that under other circumstances, indirect exploitative competition can also lead to the stronger species winning. Note that in this work, the terms “strong type” and “weak type” are used only to describe direct competitive interactions between individuals. With this understanding of the strength and weakness of competing species, the mechanism of exclusion of a strong species appears to be a paradox, since it directly contradicts the known definitions of the principle of competitive exclusion [18, 20–22]. The mechanism of indirect exploitative competition is well known to ecologists [23–25]. However, the mechanism of inverted competitive exclusion is rather impossible to model by traditional modelling methods of dependencies between the densities of individuals of competing species. This demonstrates the unprecedented effectiveness of the causal individual-based modeling we used. We have found the inverted competitive exclusion only with moderate reproduction modeled by the tripod neighborhood. In all cases studied with modeling of aggressive reproduction, the strong species won. The weak species could only win under moderate reproduction conditions. This was due to the fact that individuals of the strong species did not claim all vacancies. As a result of the moderate reproduction, there were gaps that individuals of the weak species could occupy without a direct conflict of interest. The gaps arose as a result of tripod reproduction of individuals on a hexagonal field. These free resources are the “gateway” for the entry of the offspring of a recessive species. Through strategic positioning, individuals of the recessive species reproduce in vacant microhabitats just before individuals of the dominant species attempt to occupy those microhabitats in the next iteration. This can be clearly seen at iterations 1 and 2 in Fig. 5C and at iterations 0 and 1 in Fig. 6С. This strategy allows individuals of the weak Species 2 to avoid direct competition with the strong Species 1 and deprive the dominant Species 1 of limiting reproductive resources in advance. The realization of the inverted competitive exclusion of this strategy depends not only on the ground rules, but also on the field size and the initial placement of individuals.

Inverted competitive exclusion was not realized in any of the 1000 trials at moderate reproduction on the fields of 31×31 (Fig. 4E) and 32×32 (Fig. 4F) cells under periodic boundary conditions. The fixed boundary conditions with a boundary are more favorable for the realization of this mechanism. This is because the fixed boundary of the habitat helps the weaker species survive, as the stronger species cannot attack it from the boundary.

An additional finding is that in the 31×31 field (Fig. 4E), competitors coexisted in all 1000 trials, while in the 32×32 field (Fig. 4F) competitors coexisted in 781 cases out of the 1000 cases.

The indirect exploitative exclusion mechanism of weak Species 2 is presented cell-by-cell under fixed (Fig. 5B) and periodic (Fig. 6B) boundary conditions in a 3×3 habitat model.

In the case of moderate reproduction, the possibility of indirect exploitative competition arises, which underlies the mechanism of inverted competitive exclusion. We have found that inverted competitive exclusion is realized in 33% cases (990 of 3000 trials) with moderate reproduction of competitors in models with fixed boundary conditions (Fig. 4A-C). In similar trials but with periodic boundary conditions there were 11.3% cases (339 of 3000 trials) of inverted competitive exclusion and 59.3(6)% cases (1781 of 3000 trials) of competitive coexistence were found (Fig. 4D-F).

Spatial and temporal heterogeneity of the environment is an important circumstance that contributes to the realization of competitive coexistence [26–29]. The initial habitat in our model is completely homogeneous prior to colonization by individuals. However, during the competitive colonization process, spatial heterogeneities emerge as a result of competitors’ life history activities. These heterogeneities are formed during the colonization process as the spatial distribution of five different states of the microhabitat model. These functionally emerging ecosystem spatial heterogeneities contribute to the studied mechanisms of competitive coexistence and inverted competitive exclusion.

### 3.1. Analogies of indirect exploitative competition outside the field of ecology

In indirect exploitative competition, individuals of one of the competing species, avoiding a direct conflict of interest with individuals of another species, obtain a limiting resource before individuals of the competitor, thereby depriving the latter of the opportunity to survive and/or reproduce. Our idealized model, due to its abstractness and generality, is applicable to a wide range of competition cases.

When discussing military conflicts, the concepts of asymmetric conflict are often mentioned [30–32]. These concepts include the mechanism we have modeled here, “the weak defeats the strong”, when a weak adversary, avoiding a direct confrontation, harms the strong one by depriving it of resources for military operations. The direct analogy of our logical cellular-automatic models to military models is admissible because our models reproduce the confrontation of collectives of abstract agents, which may be of different natures. The ecosystem view is the inherent context of any agent’s behavior.

### 3.2. On the definitions of the competitive exclusion principle

We investigated population dynamics under different initial conditions of the model. According to most formulations of the competitive exclusion principle, we should not have found cases of coexistence because “complete competitors cannot coexist”[20]. We found no cases of coexistence when two aggressively reproducing species colonized a large two-dimensional habitat starting from single individuals (Fig. 3). The more competitive Species 1 always displaced Species 2 as a result of the collision of their population waves (Movie S1). However, in the case of moderate reproduction under the same conditions, there were many cases of coexistence (Fig. 4), as well as a victory of the less competitive Species 2. These cases particularly interested us (Movies S2-S4).

Transparent individual-based modeling of direct and indirect competition mechanisms provides a new view of our earlier proposed formulations of the competitive exclusion principle given in the introduction. In its most succinct form, the principle of competitive exclusion states that “complete competitors cannot coexist”[20]. We formulate the classical formulation of the principle of competitive exclusion as follows:

In a spatially homogeneous, limited ecosystem, two species with different levels of competitiveness cannot coexist if they consume the same limited resource.

This formulation summarizes more than 12 well-known classical formulations of the competitive exclusion principle [18]. We have used here the important condition of other things being equal to exclude a variety of factors contributing to competitive coexistence, such as lack of any trade-offs and cooperative interactions between the competing species; reproduction of the competing species occurs only vegetatively and the species are genetically homogenous and stable; individuals of one and the same species always win individuals of other competing species in direct conflict of interest; the habitat is closed for immigration, emigration, predation, herbivory, parasitism and other disturbances; competing species are per capita identical and constant in ontogeny, in fecundity rates, in regeneration features of a microhabitat and in environmental requirements.

The mechanism of competitive coexistence that we have discovered allows species to coexist even under such conditions. This means that the formulation of the principle of competitive exclusion should be even stricter. Earlier we gave two such more stringent formulations:

*Definition 1:* “If each and every individual of a less fit species in any attempt to use any limiting resource always has a direct conflict of interest with an individual of a most fittest species and always loses, then, all other things being equal for all individuals of the competing species, these species cannot coexist indefinitely and the less fit species will be excluded from the habitat in the long run” [6].

*Definition 2:* “If a competitor completely prevents any use of at least one necessary resource by all its competitors, and itself always has access to all necessary resources and the ability to use them, then, all other things being equal, all its competitors will be excluded” [8].

The Definition 1 considers cases of direct competition for two competitors when only direct competitive relationships are realized. The Definition 2 takes into account cases of direct and indirect competition between an arbitrary number of competitors. In both cases, the conditions under which competitors cannot coexist are formulated.

The second definition of the principle has a universal application because it includes cases of both direct and indirect competition. It was formulated based on the results of consideration of the role of humans in ecological systems. Humans are the most universal consumers and are directly or indirectly responsible for the current Global Species Extinction [33–37]. Current extinction rates are about 1000 times the likely background extinction rate [38, 39]. The second definition of the principle allows us to understand the threat to biodiversity posed by the uncontrolled consumption of limited resources.

The first, earlier definition does not envision cases where a weak species can defeat a strong species through the competitive mechanism of indirect “exploitation” demonstrated in this paper. Definition 1 is centered on the condition of exclusion of a weak species through the mechanism of the direct competitive “interference”. Unlike the first our definition of the principle, the second definition lacks a focus on the competitive mechanism of “interference”. The direct conflict is not mentioned. The peculiarity of the second formulation is the possibility of multiple competitors being excluded by a single competitor. The case of exclusion of one of two competitors is also included in the formulation. The second formulation includes not only the mechanisms of direct “interference” and indirect “exploitative” but also “apparent” competition [40] because the possibility of confrontation with a shared predator, parasite or infection can be considered a special type of survival limiting resource “Apparent” and “exploitative” are two forms of indirect competition.

### 3.3. Limitations of our modeling approach

The causal individual-based mechanisms found are fully transparent due to a logical deterministic cellular automaton whose rules are based on the axioms of general ecosystem theory. These rules are relatively simple and have a causal ecological and physical interpretation [7, 8]. We believe that such transparent mathematical modeling is the best way to investigate the mechanisms of complex systems. The main limitation of using cellular automata to create transparent mathematical models is that an axiomatic theory is required for the corresponding subject domain. The culture of creating such theories has been largely lost in modern science due to the prioritization of experimental data, black box type mathematical models and opaque generative artificial intelligence [41]. Any theory, including mathematics itself, has at its core axioms from which we derive further new knowledge using logic. The complexity of creating and developing axiomatic theories has held back the development of transparent cellular-automaton modeling, which only seems relatively simple. Transparent cellular-automaton modeling is particularly promising for the development of transparent artificial intelligence systems. We recently published two articles discussing this approach in greater detail: a “Manifesto for Transparent Mathematical Modeling” [4] and “Towards eXplicitly eXplanable Artificial Intelligence” [5].

## 4. Conclusion

The ability to conduct a detailed, individual-based, causal study of the role of direct and indirect competitive relationships allowed us to identify a wide range of detailed mechanisms of competition. Complex systems consist of elements that interact with each other. Therefore, to study the mechanisms of such systems, it is very important to model them discretely and to take into account local interactions of the elements. Here we have used the glass-box artificial intelligence based on the cellular automata for individual-based modeling of direct and indirect competition. The influence of aggressive and moderate reproduction on the population dynamics of two species competing for one limiting resource, which differed only in competitiveness, has been studied. It was shown that in the case of moderate reproduction, their coexistence and even the victory of the weaker species is possible. These special cases and their mechanisms are demonstrated in videos of the computer experiments (Videos 2 and 4). To improve reproducibility, we provide the “Overview, Design Concepts, and Details” (ODD) protocol for our individual-based competition model (Supplementary 1). We also provide the complete C++ source code for the program, along with descriptions and comments (Supplementary 2). The mechanism of inverted competitive exclusion and the mechanism of coexistence of complete competitors are based on indirect exploitative competition. It is concluded that of the two formulations of the principle of competitive exclusion, which we adjusted earlier, only one takes into account not only the mechanisms of direct (interference) competition, but also the mechanisms of indirect resource competition. The results obtained are novel for theoretical ecology because they are causal and individual-based. Our modeling approach opens up prospects for mechanistic modeling of ecosystems with explicit causal consideration of local relationships between individuals of interacting species. The presented causal individual-based approach demonstrates unique capabilities for modeling complex systems. Logical-causal mathematical models open new opportunities for achieving clarity and rigor in scientific research and engineering design.

## Acknowledgments

The work has been performed under the state tasks of the Institute of Cell Biophysics of the Russian Academy of Sciences (No. 075-00607-25-00) and of the Institute of Theoretical and Experimental Biophysics of the Russian Academy of Sciences (No. 075-00223-25-03).

## Supplemental Information

**Supplementary 1.** Supplementary Overview, Design concepts and Details (ODD protocol) of the individual-based model of an ecosystem with resource competition between two species.

**Supplementary 2.** A source code of the model with Monte Carlo experiment in C++.

**Movie S1.** The strong defeats the weak. This is a classic case of competitive exclusion - the strong Species 1 defeats the weak Species 2. The habitat consists of 30×30 microhabitats with periodic boundary conditions. The mechanism of competitive exclusion is realized under aggressive propagation which is modelled by the hexagonal neighborhood.

**Movie S2.** Inverted competitive exclusion - the weak defeats the strong. A mechanism how the weak Species 2 defeats the strong Species 1. This inverted competitive exclusion is realized in a 30×30 habitat with periodic conditions under the same moderate propagation of the both competitors which is modelled by the tripod neighborhood.

**Movie S3.** The strong defeats the weak under moderate propagation. The habitat consists of 30×30 microhabitats with periodic boundary conditions. The mechanism of competitive exclusion is realized under moderate propagation which is modelled by the tripod neighborhood.

**Movie S4.** Stable coexistence of complete competitors. The habitat consists of 31×31 microhabitats with periodic boundary conditions. The mechanism of competitive coexistence is realized under moderate propagation which is modelled by the tripod neighborhood.

## References

[1] Kalmykov VL, Kalmykov LV. On ecological modelling problems in the context of resolving the biodiversity paradox. Ecological Modelling. 2016;329:1–4. doi:10.1016/j.ecolmodel.2016.03.005

[2] Tilman D. The importance of the mechanisms of interspecific competition. The American Naturalist 1987;129:769–74. 10.1086/284672

[3] Grimm V, Railsback SF. Individual-based Modeling and Ecology. STU - Student edition ed: Princeton University Press; 2005.

[4] Kalmykov VL, Kalmykov LV. Manifesto for Transparent Mathematical Modeling: from Ecology to General Science. Academia Biology. 2024;2. 10.20935/AcadBiol6166

[5] Kalmykov VL, Kalmykov LV. Towards eXplicitly eXplainable Artificial Intelligence. Information Fusion. 2025;123:103352. 10.1016/j.inffus.2025.103352

[6] Kalmykov LV, Kalmykov VL. Verification and reformulation of the competitive exclusion principle. Chaos, Solitons & Fractals. 2013;56:124–31. doi:10.1016/j.chaos.2013.07.006

[7] Kalmykov LV, Kalmykov VL. A solution to the dilemma ‘limiting similarity vs. limiting dissimilarity’ by a method of transparent artificial intelligence. Chaos, Solitons & Fractals. 2021;146:110814. 10.1016/j.chaos.2021.110814

[8] Kalmykov LV, Kalmykov VL. A Solution to the Biodiversity Paradox by Logical Deterministic Cellular Automata. Acta Biotheoretica. 2015;63:203–21. doi:10.1007/s10441-015-9257-9

[9] Kalmykov LV, Kalmykov VL. Investigation of individual-based mechanisms of single-species population dynamics by logical deterministic cellular automata. Computer Research and Modeling. 2015;7:1279–93. (in Russian). doi:10.20537/2076-7633-2015-7-6-1279-1293

[10] Kalmykov LV, Kalmykov VL. A white-box model of S-shaped and double S-shaped single-species population growth. PeerJ. 2015;3:e948. doi: 10.7717/peerj.948

[11] Park SH, Ha S, Kim JK. A general model-based causal inference method overcomes the curse of synchrony and indirect effect. Nature Communications. 2023;14:4287. doi: 10.1038/s41467-023-39983-4

[12] Nock CA, Vogt RJ, Beisner BE. Functional Traits. eLS2016. p. 1-8.

[13] Violle C, Navas M-L, Vile D, Kazakou E, Fortunel C, Hummel I, et al. Let the concept of trait be functional! Oikos. 2007;116:882–92. doi: 10.1111/j.0030-1299.2007.15559.x

[14] Gomes MP, Garcia QS, Barreto LC, Pimenta LPS, Matheus MT, Figueredo CC. Allelopathy: An overview from micro-to macroscopic organisms, from cells to environments, and the perspectives in a climate-changing world. Biologia. 2017;72:113–29. doi: 10.1515/biolog-2017-0019

[15] Grubb PJ. The maintenance of species-richness in plant communities: the importance of the regeneration niche. Biological Reviews. 1977;52:107–45. 10.1111/j.1469-185X.1977.tb01347.x

[16] Watt AS. Pattern and Process in the Plant Community. Journal of Ecology. 1947;35:1–22. doi: 10.2307/2256497

[17] Hubbell SP. Neutral theory and the evolution of ecological equivalence. Ecology. 2006;87:1387–98. doi: 10.1890/0012-9658

[18] Palmer MW. Variation in Species Richness - Towards a Unification of Hypotheses. Folia Geobotanica & Phytotaxonomica. 1994;29:511–30. doi: 10.1007/BF02883148

[19] Tilman D. Niche tradeoffs, neutrality, and community structure: A stochastic theory of resource competition, invasion, and community assembly. Proceedings of the National Academy of Sciences. 2004;101:10854–61. URL:https://www.nature.com/scitable/knowledge/library/species-interactions-and-competition-102131429/

[20] Hardin GJ. The competitive exclusion principle. Science. 1960;131:1292–7. 10.1126/science.131.3409.1292

[21] den Boer PJ. The present status of the competitive exclusion principle. Trends in Ecology & Evolution. 1986;1:25–8. 10.1016/0169-5347(86)90064-9

[22] Gause GF. The struggle for existence. Baltimore: Williams & Wilkins; 1934.

[23] Eccard J, Fey K, Caspers B, Ylönen H. Breeding state and season affect interspecific interaction types: Indirect resource competition and direct interference. Oecologia. 2011;167:623–33. doi: 10.1007/s00442-011-2008-y

[24] Lang JMB, M. E. Species Interactions and Competition. Nature Education Knowledge 2013;4:8. https://www.nature.com/scitable/knowledge/library/species-interactions-and-competition-102131429/

[25] Menge BA. Indirect Effects in Marine Rocky Intertidal Interaction Webs: Patterns and Importance. Ecological Monographs. 1995;65:21–74. 10.2307/2937158

[26] Amarasekare P. Competitive coexistence in spatially structured environments: a synthesis. Ecology Letters. 2003;6:1109–22. 10.1046/j.1461-0248.2003.00530.x

[27] Chesson P. Mechanisms of Maintenance of Species Diversity. Annual Review of Ecology and Systematics. 2000;31:343–66. doi: 10.1146/annurev.ecolsys.31.1.343

[28] Herberich MM, Gayler S, Tielbörger K. Environmental heterogeneity promotes coexistence among plant life-history strategies through stabilizing mechanisms in space and time. Basic and Applied Ecology. 2023;71:45–56. 10.1016/j.baae.2023.05.001

[29] Tamme R, Hiiesalu I, Laanisto L, Szava-Kovats R, Pärtel M. Environmental heterogeneity, species diversity and co-existence at different spatial scales. Journal of Vegetation Science. 2010;21:796–801. 10.1111/j.1654-1103.2010.01185.x

[30] Arreguín-Toft I. How the Weak Win Wars: A Theory of Asymmetric Conflict. International Security. 2001;26:93–128. doi: 10.1162/016228801753212868

[31] Pfaff CA. Anticipation in Asymmetric Warfare. In: Poli R, editor. Handbook of Anticipation: Theoretical and Applied Aspects of the Use of Future in Decision Making. Cham: Springer International Publishing; 2019. p. 1479–503.

[32] Shaohua Y. How Can Weak Powers Win? The Chinese Journal of International Politics. 2009;2:335–71. doi: 10.1093/cjip/pop004

[33] Alroy J. A multispecies overkill simulation of the end-Pleistocene megafaunal mass extinction. Science. 2001;292:1893–6. doi: 10.1126/science.1059342

[34] Branch TA, Lobo AS, Purcell SW. Opportunistic exploitation: an overlooked pathway to extinction. Trends in Ecology & Evolution. 2013;28:409–13. 10.1016/j.tree.2013.03.003

[35] Cincotta RP, Wisnewski J, Engelman R. Human population in the biodiversity hotspots. Nature. 2000;404:990–2. doi: 10.1038/35010105

[36] Diamond JM. The present, past and future of human-caused extinctions. Philosophical Transactions of the Royal Society of London B, Biological Sciences. 1989;325:469–77. 10.1098/rstb.1989.0100

[37] Vitousek PM, Mooney HA, Lubchenco J, Melillo JM. Human Domination of Earth’s Ecosystems. Science. 1997;277:494–9. 10.1126/science.277.5325.494

[38] De Vos JM, Joppa LN, Gittleman JL, Stephens PR, Pimm SL. Estimating the normal background rate of species extinction. Conservation Biology. 2015;29:452–62. 10.1111/cobi.12380

[39] Pimm SL, Jenkins CN, Abell R, Brooks TM, Gittleman JL, Joppa LN, et al. The biodiversity of species and their rates of extinction, distribution, and protection. Science. 2014;344:1246752. doi: 10.1126/science.1246752

[40] Holt RD, Bonsall MB. Apparent Competition. Annual Review of Ecology, Evolution, and Systematics. 2017;48:447–71. 10.1146/annurev-ecolsys-110316-022628

[41] Mansoulié B. Physics In Crisis: From Multiverses To Fake News: World Scientific Europe Ltd; 2022.

