## Supplementary 1 for "Individual-based modeling of direct and indirect competition by the glass-box artificial intelligence"

### Overview, Design concepts and Details (ODD protocol) of the individual-based model of an ecosystem with resource competition between two species

#### for the article "Individual-based modeling of direct and indirect competition by the glass-box artificial intelligence"

Vyacheslav L. Kalmykov[1,*], Lev V. Kalmykov[2]

[1] Laboratory of cellular stress problems, Institute of Cell Biophysics of the Russian Academy of Sciences, Pushchino, Moscow Region, Russia
[2] Laboratory of cell engineering, Institute of Theoretical and Experimental Biophysics of the Russian Academy of Sciences, Pushchino, Moscow Region, Russia

* Corresponding author.

ODD is a standard protocol for describing individual-based and agent-based models. The main goals of ODD are to make model descriptions more understandable, complete, and efficient, to improve the rigorous formulation of models, and to make the models more reproducible (Grimm et al. 2006; Grimm et al. 2010; Grimm et al. 2020). The methodology of our modeling approach has been presented previously (Kalmykov & Kalmykov 2013; Kalmykov & Kalmykov 2015a; Kalmykov & Kalmykov 2015b; Kalmykov & Kalmykov 2021; Kalmykov & Kalmykov 2024), but the ODD protocol of our ecosystem model is presented here for the first time. We describe the ecosystem model as a cellular automaton: we define a cellular automaton field with a type of boundary conditions, a set of possible states of each cell in the field, a neighborhood of local interactions, a function of transitions between states for each cell in the field, and an initial pattern. We defined all these entities and gave a biological interpretation of the model. The deterministic logic cellular automaton models causal mechanisms of interspecific competition based on local agents. The complete source code of the model with detailed comments (352 lines of code) is provided in the "Supplementary II Source Code" file for reproducing the model. Information on the compiler and code libraries is provided.

#### ODD protocol for our individual-based model of ecosystem

#### A. Overview

##### 1. Purpose:

The overall goal of our model is to investigate individual-based mechanisms of a new paradoxical mechanism of indirect exploitative competition - the weak competitor can defeat the strong. Individual-based models are synonymous with agent-based models (Vincenot 2018). Their main specificity is the direct possibility to take into account local causal interactions of their micro-objects. This is the main essence of this mechanistic approach. The micro-objects of our model are micro-ecosystems corresponding to the cells of a cellular-automaton field. An individual microecosystem can be a free microhabitat containing life history resources for one individual, or a microhabitat inhabited by an individual, or a habitat in the regime of resource regeneration after the death of an individual, or even a microhabitat without resources, which in principle is unsuitable for providing life history resources for individuals of a given species (e.g., a stone in the case of the lawn grass model). The rules of the cellular automaton determine all possible changes in the states of each and every microecosystem. The cellular-automaton neighborhood defines the semantics of the local environment of each microecosystem when implementing cellular automaton rules. The rules of the cellular automaton underlying our models are formulated on the basis of the logical axioms of ecosystem theory.

From a physical point of view, we model and investigate mechanisms of propagation and collision of population waves of two types propagating in a homogeneous bounded environment with a renewable resource. The hypothesis we test is whether it is possible for a weaker species to win in competition with a stronger one, other things being equal, under conditions of moderate reproduction of the both species on a two-dimensional field. The biological analog is an ecosystem with two species of lawn grasses that reproduce only vegetatively by shoots. The reproduction is modeled using a tripod neighborhood. Competition manifests itself in localized conflicts of interest over microhabitats with a resource. The offspring of the more competitive species occupies the disputed microhabitat. Individuals of competing species are identical in all characteristics except competitiveness. We define competitiveness as the value that determines the outcome of competition under direct simultaneous conflict of interest between individuals for a single unit of a single limiting resource. Such a single unit of limiting resource in our models is one free microhabitat containing all resources necessary for the life activity of one offspring of any individual of competing species. The model conditions allow us to rigorously and transparently test our hypothesis that a weak species can defeat a strong one given other equal characteristics of both species. In doing so, the subject individual-based mechanisms are clearly revealed.

**2. Entities, state variables, and scales:**

Our customized model is a logical deterministic cellular automaton. To complete characterize such a model, we provide a description of the field with boundary conditions, the set of possible states of each cell of the field, the neighborhood of the cellular automaton, the transition function between the states of each cell of the field, and the initial pattern from which the evolution of the model begins.

We use a one-dimensional (Fig. 1) and a two-dimensional cellular automaton field (Fig. 3-6 and Movies S1-S3). We study periodic conditions of two types: (i) when the field has a boundary and (ii) when it is closed by periodic conditions to avoid edge effects. The cellular automaton field consisting of hexagonal cells has the shape of a parallelogram. Periodic boundary conditions connect the boundaries of the field, i.e. a torus is obtained.

The set of possible states of the field cell is {0, 1, 2, 3, 4, 5}:

0 - a free microhabitat which can be occupied by an individual of both species;

1 - a microhabitat is occupied by a living individual of the first species;

2 - a regeneration state of a microhabitat after death of an individual of the first species;

3 - a microhabitat is occupied by a living individual of the second species;

4 - a regeneration state of a microhabitat after death of an individual of the second species;

5 - a boundary state of a field cell. In case of periodic boundary conditions and homogeneous habitat this state is not used.

Aggressive vegetative reproduction is modeled by the hexagonal neighborhood of the cellular automaton (Fig. 2A), and moderate reproduction is modeled by the tripod neighborhood (Fig. 2B).

The function f1 of transitions between states of each cell of the field contains logical deterministic rules (Fig. 2C):

#### B. Design concepts

**1. Process overview and scheduling**

Each state of a field cell lasts for one iteration of the cellular automaton. The function f1 of transitions between the states of each cell of the field contains logical deterministic rules (Fig. 2):
0→0, a microhabitat remains free if there is no one living individual in the neighborhood;
0→1, a microhabitat will be occupied by an individual of the first species if there is at least one individual of the first species in its neighborhood (priority over the second species);
0→3, a microhabitat will be occupied by an individual of the second species if there is at least one individual of the second species in its neighborhood and there is no one individual of the first species in its neighborhood;
1→2, after death of an individual of the first species its microhabitat goes into the regeneration state;
2→0, after the regeneration state a microhabitat will be free if there is no one living individual in the neighborhood;
2→1, after the regeneration state a microhabitat will be occupied by an individual of the first species if there is at least one individual of the first species in its neighborhood (priority over the second species);
2→3, after the regeneration state a microhabitat will be occupied by an individual of the second species if there is at least one individual of the second species in its neighborhood and there is no one individual of the first species in its neighborhood;
3→4, after death of an individual of the second species its microhabitat goes into the regeneration state;
4→0, after the regeneration state a microhabitat will be free if there is no one living individual in its neighborhood;
4→1, after the regeneration state a microhabitat will be occupied by an individual of the first species if there is at least one individual of the first species in its neighborhood (priority over the second species);
4→3, after the regeneration state a microhabitat will be occupied by an individual of the second species if there is at least one individual of the second species in its neighborhood and there is no one individual of the first species in its neighborhood;
5→5, a boundary state of a field cell passes into itself.

Based on these rules, automatic individual- based logic inference is realized at the level of each cell of the field and the neighborhood of each cell of the field. Thus, on the basis of the micro-level of each cell and mini-levels of each neighborhood all states of all cells of the field are determined, i.e. macro-level. The cellular automaton neighborhood and the rules of microhabitat state transitions provide the semantics of gluing the state dynamics of all microhabitats into the overall state dynamics of the ecosystem as a whole.

#### C. Details

##### 1. Initialization

Our model is completely deterministic, so initial conditions can have a significant impact on the model evolution. Such initial conditions are the size of the field, the boundary conditions of the field, and the states of the field cells at the initial iteration (initial pattern).

The initial pattern consists of a bounded homogeneous habitat in which we place one individual of the first species and one individual of the second species. This placement can be different and lead to different population dynamics. We investigate this by automating the process of placing individuals in the initial iteration in a probabilistic manner. All subsequent dynamics of the model are complete deterministic. We run a series of 1000 such experiments. This constitutes a Monte Carlo experiment (n=1000).

##### 2. Input data

Black-box phenomenological models can accurately fit experimental data, but they do not model discrete elements and their local interactions. Mechanistic white box models model discrete elements and their local interactions. The elementary object of the model, the states it can be in and logical transitions between these states are defined on the basis of our theory of the subject area. Input data in our model is a combination of field type and size, type of boundary conditions, type of cellular automaton neighborhood, cellular automaton rules, and the initial pattern of states of the field cells.

##### 3. Submodels:

The model has three levels of organization - micro, mini and macro.

The micro level is each individual cell of the field.

The mini level is the cellular automaton neighborhood of each cell of the field.

The macro level is the entire field.

The model is relatively simple, but can be extended indefinitely with additional number of states, neighborhood types and nested fields.

Subsections of the model program are commented in detail in its source-code.

#### References


Grimm V, Berger U, Bastiansen F, Eliassen S, Ginot V, Giske J, Goss-Custard J, Grand T, Heinz S, Huse G, Huth A, Jepsen J, Jørgensen C, Mooij W, Müller B, Pe'er G, Piou C, Railsback S, Robbins A, and Deangelis D. 2006. A Standard Protocol for Describing Individual-Based and Agent Based Models. *Ecological Modelling* 198:115-126. 10.1016/j.ecolmodel.2006.04.023

Grimm V, Berger U, DeAngelis DL, Polhill JG, Giske J, and Railsback SF. 2010. The ODD protocol: A review and first update. *Ecological Modelling* 221:2760-2768. https://doi.org/10.1016/j.ecolmodel.2010.08.019

Grimm V, Railsback SF, Vincenot CE, Berger U, Gallagher C, DeAngelis DL, Edmonds B, Ge J, Giske J, Groeneveld J, uuml, rgen, Johnston ASA, Milles A, Nabe-Nielsen J, Polhill JG, Radchuk V, Rohw, auml, der M-S, Stillman RA, Thiele JC, Ayll, oacute, and n D. 2020. The ODD Protocol for Describing Agent-Based and Other Simulation Models: A Second Update to Improve Clarity, Replication, and Structural Realism. *Journal of Artificial Societies and Social Simulation* 23:7. 10.18564/jasss.4259

Kalmykov LV, and Kalmykov VL. 2013. Verification and reformulation of the competitive exclusion principle. *Chaos, Solitons & Fractals* 56:124-131. http://dx.doi.org/10.1016/j.chaos.2013.07.006

Kalmykov LV, and Kalmykov VL. 2015a. A Solution to the Biodiversity Paradox by Logical Deterministic Cellular Automata. *Acta Biotheoretica* 63:203-221. http://dx.doi.org/10.1007/s10441-015-9257-9

Kalmykov LV, and Kalmykov VL. 2015b. A white-box model of S-shaped and double S-shaped single-species population growth. *PeerJ* 3:e948. http://dx.doi.org/10.7717/peerj.948

Kalmykov LV, and Kalmykov VL. 2021. A solution to the dilemma `limiting similarity vs. limiting dissimilarity' by a method of transparent artificial intelligence. *Chaos, Solitons & Fractals* 146:110814. https://doi.org/10.1016/j.chaos.2021.110814

Kalmykov VL, and Kalmykov LV. 2024. Manifesto for Transparent Mathematical Modeling: from Ecology to General Science. *Academia Biology* 2. https://doi.org/10.20935/AcadBiol6166

Vincenot CE. 2018. How new concepts become universal scientific approaches: insights from citation network analysis of agent-based complex systems science. *Proceedings of the Royal Society B: Biological Sciences* 285:20172360. doi:10.1098/rspb.2017.2360
