## Supplementary 2 for "Individual-based modeling of direct and indirect competition by the glass-box artificial intelligence"

### A source code of the model with Monte Carlo experiment in C++ for the article “Individual-based modeling of direct and indirect competition by the glass-box artificial intelligence”

Vyacheslav L. Kalmykov[1,*], Lev V. Kalmykov[2]

[1] Laboratory of cellular stress problems, Institute of Cell Biophysics of the Russian Academy of Sciences, Pushchino, Moscow Region, Russia
[2] Laboratory of cell engineering, Institute of Theoretical and Experimental Biophysics of the Russian Academy of Sciences, Pushchino, Moscow Region, Russia

* Corresponding author.

This source code is an extension of the previously published code of the basic model of an ecosystem with interspecific competition for resources (Kalmykov & Kalmykov 2021; Kalmykov & Kalmykov 2024). A Monte Carlo simulation code was added to study variations in initial conditions. A hexagonal cellular automaton neighborhood was used to model aggressive reproduction, and a tripod neighborhood was used to model moderate reproduction. To study the role of boundary conditions, periodic boundary conditions were supplemented with fixed field boundaries.

The model is a deterministic individual-based cellular automaton and implements automatic deductive inference based on the logical rules of our axiomatic-deductive ecosystem theory (Kalmykov & Kalmykov 2015; Kalmykov & Kalmykov 2021). The source code of the model is written in C++ programming language and tested in Microsoft Visual C++ Studio 2015.

The advantage of our program is the complete transparency of all local events of the simulated system. The presented code demonstrates exactly such a transparent model. The evolution of the system occurs iteratively. At each iteration, all elements of the system are displayed.

To investigation patterns at each iteration, it is necessary to use the function *_getch()*. The basic logical rules for automatic inference are contained in the function *f1*. Each element of the modelled ecosystem is clearly visualized. This is a method of transparent artificial intelligence, which made it possible to test the validity of the mutually exclusive hypotheses of limiting similarity and limiting dissimilarity. The parameters of the competitiveness of the species are set as follows. The field size can be changed by varying the parameter “l_size”. The number of individuals of the Species 1 is output to the file "data_1.txt" at each iteration. The number of individuals of the Species 2 is output to the file "data_2.txt" at each iteration. The file "data_3.txt" save numbers when Species 1 wins, Species 2 wins, Species 1 and Species 1 coexist and when both competitors extinct.

In the specified code listing the field size is 34x34. For visualization, you may need to increase the size of the application window and reduce the font in the properties of the console application. The initial pattern of the habitat is populated by single individuals of two competing species.

The size of the habitat is specified by the l_size parameter.

In this program, the boundary conditions are non-periodic, and the boundary is modeled by the cell’s state “5”, which cannot be changed by iterations of the cellular automaton. To model periodic conditions, we need to use the *Torus* function (Kalmykov & Kalmykov 2021; Kalmykov & Kalmykov 2024):

```
int Torus(int x)
{
	if (x<0) return x + l_size;
	else return x % l_size;
}
```

**Table 1. Description of variables and symbols of the source code of the model**

| Symbol | Description |
|---|---|
| l_size | This variable defines the array length. The field size calculates as l_size *l_size |
| s | This is a central cell of the neighborhood. It is defined by the array element with indices (i, j), where i and j are integer numbers |
| s1 | This is a cell of the neighborhood. It is defined by the array element with indices (i-1, j+1), where i and j are integer numbers |
| s2 | This is a cell of the neighborhood. It is defined by the array element with indices (i+1, j), where i and j are integer numbers |

| | |
|---|---|
| s3 | This is a cell of the neighborhood. It is defined by the array element with indices (i, j-1), where i and j are integer numbers |
| sp1_wins | The number how many times Species 1 wins |
| sp2_wins | The number how many times Species 2 wins |
| sp1_sp2_coexist | The number how many times both Species coexist |
| sp1_sp2_die | The number how many times both Species extinct |
| iteration | The number of iteration |
| experiment_number | The number of Monte Carlo experiment |
| a | The number of individuals of the Species 1 |
| b | The number of individuals of the Species 2 |
| aa | Final number of individuals of the Species 1 |
| bb | Final number of individuals of the Species 2 |
| t1 | First index of an element- initial position of an individual of the Species 1 |
| t2 | Second index of an element - initial position of an individual of the Species 1 |
| t3 | First index of an element- initial position of an individual of the Species 2 |
| t4 | Second index of an element - initial position of an individual of the Species 2 |

**References**


Kalmykov LV, and Kalmykov VL. 2015. A Solution to the Biodiversity Paradox by Logical Deterministic Cellular Automata. *Acta Biotheoretica* 63:203-221. http://dx.doi.org/10.1007/s10441-015-9257-9

Kalmykov LV, and Kalmykov VL. 2021. A solution to the dilemma `limiting similarity vs. limiting dissimilarity' by a method of transparent artificial intelligence. *Chaos, Solitons & Fractals* 146:110814. https://doi.org/10.1016/j.chaos.2021.110814

Kalmykov VL, and Kalmykov LV. 2024. Manifesto for transparent mathematical modeling: from ecology to general science. *Academia Biology* 2:1-9. 10.20935/AcadBiol6166 https://doi.org/10.20935/AcadBiol6166

```
#include "stdafx.h"
#include <iostream>
#include <conio.h>
#include <stdlib.h>
#include <time.h>
#include <windows.h>
#include <fstream>

using namespace std;

const char * file1 = "data_1.txt"; /* File with the number of
individuals of the Species 1 */
const char * file2 = "data_2.txt"; /* File with the number of
individuals of the Species 2 */

const char * file3 = "data_3.txt";// Statistics

const int l_size = 34; /* The field size (ecosystem) is of 34x34
cells (microhabitats), l_size>3 */

double random()
{
	return rand() / (double(RAND_MAX));
}

int c[l_size][l_size]; // Main array
int c_temp[l_size][l_size]; // Temporary array

/* f1 is the function of transitions between the states of a field
cell.
The function contains rules for automatic logical inference.
The arguments of the f1 function are the states of cells of the
tripod neighborhood. */
int f1(int s, int s1, int s2, int s3)
{
	if (s == 5) return 5; // Boundary

	/* Rules of propagation of individuals of the Species 1.
	The condition for the Species 1 is verified before the
condition for the Species 2.
	A free microhabitat and a microhabitat after the regeneration
state may be occupied by an individual of the Species 1, if there is
at least one individual of the Species 1 in the neighborhood. */
	if (((s1 == 1) || (s2 == 1) || (s3 == 1)) &&
		((s == 0) || (s == 2) || (s == 4)))
		return 1;

	/* Rules of propagation of individuals of the Species 2.
	The condition for the Species 2 is verified after the
condition for the Species 1.
	A free microhabitat and a microhabitat after the regeneration
state may be occupied by an individual of the Species 2, if there is
at least one individual of the Species 2 in the neighborhood. */
	if (((s1 == 3) || (s2 == 3) || (s3 == 3)) &&
		((s == 0) || (s == 2) || (s == 4)))
		return 3;

	/* After the death of an individual of the Species 1 its
microhabitat goes into the state of resource regeneration */
	if (s == 1) return 2;
	/* After the regeneration state a microhabitat becomes free
if there is no a single individual in its neighborhood */
```

```
	if (s == 2) return 0;

	/* After the death of an individual of the Species 2 its 
microhabitat goes into the state of resource regeneration */
	if (s == 3) return 4;
	/* After the regeneration state a microhabitat becomes free 
if there is no a single individual in its neighborhood */
	if (s == 4) return 0;

	/* A microhabitat remains free if there is no a single 
individual in its neighborhood */
	if (s == 0) return 0;
}

int main()
{
	system("Color F0"); // Field color - white
	srand((unsigned)time(NULL));

	int sp1_wins = 0;
	int sp2_wins = 0;
	int sp1_sp2_coexist = 0;
	int sp1_sp2_die = 0;


	// Monte Carlo simulation with n=1000 experiments
	for (int experiment_number = 1; experiment_number <= 1000; 
experiment_number++)
	{
		int aa = 0;
		int bb = 0;

		int iteration = 0; // Initial iteration

							// Creation of the 
initial pattern
		for (int i = 0; i<l_size; i++)
			for (int j = 0; j<l_size; j++)
			{
				c[i][j] = 0;
				c_temp[i][j] = 0;
			}


		for (int i = 0; i<l_size; i++)
		{
			c[i][0] = 5;
			c[i][l_size - 1] = 5;
		}
		for (int j = 0; j<l_size; j++)
		{
			c[0][j] = 5;
			c[l_size - 1][j] = 5;
		}


		// Random initial placement of individuals on the 
field
		int t1 = 0; int t2 = 0;	int t3 = 0; int t4 = 0;
		// Random coordinates of the position of an individual 
of the Species 1
	T1:	t1 = rand() % (l_size - 1);
```

```
		t2 = rand() % (l_size - 1);
		if ((t1 == 0) || (t2 == 0)) goto T1;
		// Random coordinates of the position of an individual 
of the Species 2
	I1: t3 = rand() % (l_size - 1);
		t4 = rand() % (l_size - 1);
		// If the coordinates match, then new coordinates are 
generated
		if ((t3 == 0) || (t4 == 0)) goto I1;
		if ((t1 == t3) && (t2 == t4)) goto I1;

		c[t1][t2] = 1; // Initial position of an individual of 
the Species 1
		c[t3][t4] = 3; // Initial position of an individual of 
the Species 2


		for (; iteration <= 100;) // Duration of one 
experiment
		{

			// Calculation the number of individuals in the 
habitat
			int a = 0; // Number of individuals of the 
Species 1
			int b = 0; // Number of individuals of the 
Species 2
			for (int i = 0; i<l_size; i++)
				for (int j = 0; j<l_size; j++)
				{
					if (c[i][j] == 1) a++;
					if (c[i][j] == 3) b++;
				}
			// Final number of individuals of the Species 1
			if (a == 0) aa++;
			// Final number of individuals of the Species 2
			if (b == 0) bb++;


			// Console visualization
			cout << "\n";


			for (int i = 0; i<l_size; i++)
			{
				for (int k = 0; k <= i; k++)
					cout << " ";
				for (int j = 0; j<l_size; j++)

				{
					if (c[i][j] == 0)
					{
						HANDLE hOut;
						hOut = 
GetStdHandle(STD_OUTPUT_HANDLE);

	SetConsoleTextAttribute(hOut, BACKGROUND_RED | 
BACKGROUND_GREEN | BACKGROUND_BLUE | BACKGROUND_INTENSITY | 7);
						cout << " 0";
					}

					if (c[i][j] == 1)
```

```
                                {
                                        HANDLE hOut;
                                        hOut =
GetStdHandle(STD_OUTPUT_HANDLE);

        SetConsoleTextAttribute(hOut, BACKGROUND_RED |
BACKGROUND_GREEN | BACKGROUND_BLUE | BACKGROUND_INTENSITY |
FOREGROUND_RED | FOREGROUND_INTENSITY | 0);
                                        cout << " 1";
                                }

                                if (c[i][j] == 2)
                                {
                                        HANDLE hOut;
                                        hOut =
GetStdHandle(STD_OUTPUT_HANDLE);

        SetConsoleTextAttribute(hOut, BACKGROUND_RED |
BACKGROUND_GREEN | BACKGROUND_BLUE | BACKGROUND_INTENSITY | 7);
                                        cout << " 2";
                                }

                                if (c[i][j] == 3)
                                {
                                        HANDLE hOut;
                                        hOut =
GetStdHandle(STD_OUTPUT_HANDLE);

        SetConsoleTextAttribute(hOut, BACKGROUND_RED |
BACKGROUND_GREEN | BACKGROUND_BLUE | BACKGROUND_INTENSITY |
FOREGROUND_BLUE | FOREGROUND_INTENSITY | 0);
                                        cout << " 3";
                                }

                                if (c[i][j] == 4)
                                {
                                        HANDLE hOut;
                                        hOut =
GetStdHandle(STD_OUTPUT_HANDLE);

        SetConsoleTextAttribute(hOut, BACKGROUND_RED |
BACKGROUND_GREEN | BACKGROUND_BLUE | BACKGROUND_INTENSITY | 7);
                                        cout << " 4";
                                }

                                if (c[i][j] == 5)
                                {
                                        HANDLE hOut;
                                        hOut =
GetStdHandle(STD_OUTPUT_HANDLE);

        SetConsoleTextAttribute(hOut, BACKGROUND_RED |
BACKGROUND_GREEN | BACKGROUND_BLUE | BACKGROUND_INTENSITY | 0);
                                        cout << " 5";
                                }

                        }

                        cout << "\n";
                }

                cout << "\n";
```

```
			// Output statistics
			HANDLE hOut;
			hOut = GetStdHandle(STD_OUTPUT_HANDLE);
			SetConsoleTextAttribute(hOut, BACKGROUND_RED |
BACKGROUND_GREEN | BACKGROUND_BLUE | BACKGROUND_INTENSITY);

			cout << " Number of Monte Carlo experiment: "
<< experiment_number << "\n";
			cout << " Number of iterations: " << iteration
<< "\n";
			cout << " Number of individuals of the Species
1: " << a << "\n";
			cout << " Number of individuals of the Species
2: " << b << "\n";

			/* Calculation the states of the field cells in
the next iteration.
			The arguments of the f1 function are the states
of cells of the tripod neighborhood. */
			for (int i = 0; i<l_size; i++)
				for (int j = 0; j<l_size; j++)
					c_temp[i][j] = f1(c[i][j],

							c[i - 1][j +
1],
							c[i + 1][j],

							c[i][j - 1]);

			/* Transfer of the new calculated states of the
field cells to the Main array. */
			for (int i = 0; i<l_size; i++)
				for (int j = 0; j<l_size; j++)
					c[i][j] = c_temp[i][j];


			// File with the number of individuals of the
Species 1
			if (aa<2)
			{
				ofstream fout1(file1, ios::out |
ios::app);
				fout1 << a << " ";

				if (iteration == 100) // Move to a new
line at the end of the experiment
				{
					fout1 << "\n";
					fout1.close();
				}

				else
					fout1.close();
			}

			else
			{
				ofstream fout1(file1, ios::out |
ios::app);
				fout1 << " ";
				if (iteration == 100)
```

```
                        {
                               fout1 << "\n";
                               fout1.close();
                        }
                        else
                               fout1.close();
                 }


                 // File with the number of individuals of the
Species 2
                 if (bb<2) {
                        ofstream fout2(file2, ios::out |
ios::app);
                        fout2 << b << " ";

                        if (iteration == 100) // Move to a new
line at the end of the experiment
                        {
                               fout2 << "\n";
                               fout2.close();
                        }

                        else
                               fout2.close();
                 }

                 else
                 {
                        ofstream fout2(file2, ios::out |
ios::app);
                        fout2 << " ";
                        if (iteration == 100)
                        {
                               fout2 << "\n";
                               fout2.close();
                        }
                        else
                               fout2.close();
                 }


                 // Statistics
                 if (iteration == 100)
                 {
                        {
                               if ((b == 0) && (a != 0))
sp1_wins++;
                               if ((a == 0) && (b != 0))
sp2_wins++;
                               if ((a != 0) && (b != 0))
sp1_sp2_coexist++;
                               if ((a == 0) && (b == 0))
sp1_sp2_die++;
                        }
                        if ((iteration == 100) &&
(experiment_number == 1000))
                        {
                               ofstream fout3(file3, ios::out |
ios::app);
                               fout3 << "sp1 wins: " << sp1_wins
<< "\n"
```

```
                                                << "sp2 wins: " << sp2_wins
<< "\n"
                                                << "sp1 & sp2 coexist: " <<
sp1_sp2_coexist << "\n"
                                                << "sp1 & sp2 die: " <<
sp1_sp2_die << "\n";
                                        fout3.close();
                                }
                        }


                        //_getch(); // Pause
                        iteration++; // Counter of iterations
                        system("cls"); // Clear screen
                }
        }
        return 0;
}
```
